# Protein kinase D promotes prostate cancer cell bone metastasis by positively regulating Runx2 in a MEK/ERK1/2-dependent manner

**DOI:** 10.1101/2022.04.16.488551

**Authors:** Adhiraj Roy, Sahdeo Prasad, Yapeng Chao, Jinjun Zhao, Qiming Jane Wang

## Abstract

Most cancer patients die because of tumor metastasis not due to tumors at the primary site. Prostate tumors in advance stages frequently metastasize to the bone, which is the main cause of death for the disease. The family of protein kinase D (PKD) has been implicated in prostate cancer development, however, its role in prostate cancer metastasis has not been investigated. In this study, we examined the contribution of PKD to the metastatic potential of prostate cancer cells and the impact of PKD inhibition on prostate cancer bone metastasis *in vivo*. Our data showed that depletion of PKDs by siRNA or inhibition of PKD by an inhibitor CRT0066101 in a highly invasive prostate cancer PC3-ML cells potently inhibited colony formation and cell migration. Furthermore, depletion or inhibition of PKD significantly blocked invasion of PC3-ML cells and suppressed expression of genes related to bone metastasis. The reduced invasive activity resulted from PKD depletion was in part mediated through the transcription factor Runx2 as its silencing decreased PKD-mediated metastatic gene expression. Mechanistically, our data indicate that PKD modulated Runx2 target gene expression through the MEK/ERK1/2 signaling axis. Additionally, we examined whether PKD inhibitor CRT0066101 could prevent prostate cancer bone metastasis in a mouse model of metastasis, where intracardiac injection of PC3-ML cells led to metastasis of cells to the bone. We found that CRT0066101 potently decreased the frequency of micrometastases in mouse bone. These results indicate that PKDs play an important role in bone metastasis of prostate cancer cells and its inhibition may be beneficial for treatment of advanced stages of cancer.

## Introduction

Prostate cancer is the most common cancer in American men after skin cancer. It is also one of the leading causes of cancer death among men of all races. Despite effective initial management and improved therapeutic options in recent years, approximately half of prostate cancer patients reported to die from distal metastases (1). Bone is one of the most preferred sites of metastasis of prostate cancer cells with up to 90% of patients with advanced disease having bone metastases (2). Bone metastases invariably lead to hypercalcemia, bone pain, fractures and nerve compression, increase the morbidity and mortality of prostate cancer patients (3).

PKD is a family of serine/threonine kinases that belong to a subfamily of the Ca^2+^/calmodulin-dependent protein kinase (CaMK) superfamily. Three isoforms of PKD (PKD1, 2, 3) with high sequence homology have been identified. PKD can be activated in a canonical pathway through phosphorylation of two conserved serine residues by classical or novel PKCs. PKD regulates many important cancer-associate protein targets including β-catenin, androgen receptor, Wnt5, urokinase-type plasminogen activator (uPA), Hsp27, Class IIa HDACs, VEGF, matrix metalloproteases (MMPs), and MEK/ERK, etc (4). Its important role in cancer development provides the foundation for targeting PKDs by small molecules for cancer treatment. Several small molecule inhibitors of PKD have been developed including CID755673 and its analogs kb-NB142–70, SD-208, 1-naphthyl PP1 (1-NA-PP1), Compound 139, CRT0066101 and others (5-9). Among them CRT0066101, an aminopyridine arene, emerged as a highly potent, cell-permeable, and *in vivo* active pan-PKD inhibitor. CRT0066101 was found to block tumor cell proliferation, migration and invasion and inhibit growth of pancreatic, breast, bladder, and colorectal tumor xenografts in mice (10-13).

Growing evidence supports an important role of PKD in prostate cancer (9,14). Aberrant expression of PKD has been reported in human prostate carcinoma (15,16). In particular, PKD2 and PKD3 are frequently expressed in highly metastatic prostate cancer cells and have been shown to promote prostate tumor cell proliferation and survival. PKD2 and PKD3 have also been implicated in prostate cancer cell invasion and migration by promoting NF-κB and uPA expression and increased MMP-9 expression (17-19). PKD3 increases prostate cancer cell survival by increasing prolonged activation of Akt and ERK1/2 (16). PKD3 has also been shown to promote the production of tumor-promoting factors as knockdown of PKD3 inhibited the production of MMP-9, IL-6, IL-8, and GROα from prostate cancer cells (19). Despite these known functions of PKDs, the role of PKDs in prostate cancer bone metastasis is not yet known.

The transcription factor Runx2 is involved in many cellular and physiological functions including the regulation of bone formation, bone turnover and epigenetic gene control during cell division and abnormal or high expression of Runx2 transcription factor has been associated with tumor cells that metastasize to the bone (20). In this study, we investigated whether PKD play a role in prostate cancer cell migration/invasion via modulating Runx2 expression. We further determined whether inhibition of PKD prevents bone metastasis of prostate cancer cells in a mouse model of metastasis. Using PC3-ML cells, a subline derived from PC3 cell line having 80% metastatic efficiency to the lumbar vertebrae and produces bone metastasis in mice (21), we found that inhibition of PKD by CRT0066101 potently inhibited PC3-ML cell metastasis to the bone. Silencing of PKD2/3 or inhibition of PKD in PC3-ML cells abrogated tumor cell migration and invasion in part through downregulating bone metastatic genes via reducing Runx2 protein expression. Our results identified PKD2/3-Runx2 as novel targets in bone metastasis of prostate cancer and targeted suppression of these enzymes may have significant therapeutic implication.

## Materials and Methods

### Materials

RPMI 1640 medium was purchased from Cellgro (Manassas, VA) and other culture materials were from Invitrogen (Life Technologies, Carlsbad, CA). Antibodies against PKD2, PKD3, Runx1, Runx2, and Runx3 were obtained from Cell Signaling Technology. GAPDH was purchased from Enzo Lifescience. All other chemicals were purchased from Sigma unless otherwise stated.

### Cell culture

Human prostatic tumor variant PC-3 ML cells and PC3-ML cells infected with ZsGreen Lentivirus was kindly provided by Dr. Marcelo G. Kazanietz, Perelman School of Medicine, University of Pennsylvania, Philadelphia. PC3-ML cells were cultured in RPMI 1640 medium supplemented with 10% FBS, 100 U/ml penicillin, and 100 μg/ml streptomycin.

### Western blot analysis

Western blot analysis was carried out as previously reported (22). Briefly, protein concentration was determined using Pierce BCA Protein Assay kit (Thermo Fisher Scientific, Rockford, IL) according to manufacturer’s instructions. Equal amount of total protein was resolved by SDS-polyacrylamide gel electrophoresis (SDS-PAGE). After electrophoresis, proteins were transferred onto nitrocellulose membranes. Membranes were blocked with 5% non-fat dry milk in Tris-buffered saline containing 0.2% Tween 20 for 1 h, then blotted with a primary antibody, followed by a secondary goat anti-rabbit or anti-mouse antibody conjugated to horseradish peroxidase. Blots were developed using an Enhanced Chemiluminescence (ECL) reagent (GE Healthcare).

### Matrigel Invasion Assays

Matrigel invasion assay was conducted as previously described. Briefly, cells were seeded (1.5 × 10^4^ cells/well) in RPMI medium containing 1% FBS in the upper compartment of a Boyden chamber with Matrigel-coated 12 μm pore polycarbonate membrane insert (NeuroProbe, Gaithersburg, MD). RPMI medium containing 20% FBS was used in the lower chamber. After an incubation period of 22 h at 37°C, membranes were recovered and cells on the upper side of membrane were wiped off and then stained with the DIFF Quik Stain Set (Dade Behring, Deerfield, IL). Similar procedures were done with control chamber without matrigel. Migratory cells in each well were counted by phase contrast microscopy in 5 random fields and analyzed. Percent invasion was determined by dividing the number of invaded cells by the total number of cells migrated through the control insert.

### Colonigenic assay

To determine the growth of prostate cancer cells as tumor mass, we conducted a colonigenic assay. PC3-ML (500 cells/well) was seeded in 6-well plates and allowed to adhere overnight then transfected with siRNA against PKD2 and PKD3 as well as treated with CRT0066101. Cells were allowed to form colonies for 9 days, and then stained with 0.25% crystal violet solution. Number of colonies in each well were counted and then analyzed.

### Wound healing assay

PC3-ML cells were grown to confluence in 6-well plates. Wounds were created by scraping the monolayer with a pipette tip and then wash with fresh media to remove the floating cells. Cells were transfected with PKD2 or PKD3 siRNA or exposed to CRT0066101 separately. An image of wound was captured immediately and after 24 hours under an inverted phase-contrast microscope. The wound gap was measured by NIH ImageJ software and percent wound healing was calculated.

### Animals

The 4-week-old male athymic nu/nu mice were obtained from the breeding colony of the Charles River (Wilmington, MA). The animals were housed and maintained with food, light, and water in standard manner. This animal protocol was approved by University of Pittsburgh Institutional Animal Care and User Committee (IACUC).

### Bone metastasis in nude mice

Prostate cancer PC3-ML cell lines, stably transfected with ZsGreen lentivirus, were harvested from sub-confluent culture. Cells were washed and resuspended in PBS and then kept in ice. Single cell suspension (50,000 cells/mouse) with more than 90% viability were inoculated in the left cardiac ventricle of anaesthetized athymic nude mice (n=12). During cell inoculation, needle was gently inserted straight down into the ventricle. Same day animals were randomly assigned into two experimental groups (1) vehicle (5% dextrose, oral gavage daily) and (2) CRT0066101 (80 mg/kg dissolved in 5% dextrose; oral gavage daily). Animals were sacrificed after 4 weeks by placing in a CO_2_ chamber followed by cervical dislocation. Femur bone from each animal was dissected out for further study. To confirm the delivery of cells into the systemic blood circulation, blue fluorescent 10 μm polystyrene beads (Molecular Probes, Eugene, OR) were injected separately into the left cardiac ventricle of mice in same manner. After 24 hours, animals were sacrificed and the presence of fluorescent blue beads were observed in kidney and lungs under fluorescent microscope.

### Decalcification of bone and micrometastases analysis

The cleaned collected bones were transferred to 4% paraformaldehyde solution. After 24 h bones were transferred to fresh paraformaldehyde for an additional period of 24 h. Decalcification of bone was performed by transferring the bone to a 0.5 M EDTA (pH 7.2) solution for 10 days. EDTA solution was changed every day by fresh solution and maintained at 4^0^C. Bone was placed in a rocker throughout entire decalcification process. After 10 days, bones were incubated in 30% sucrose for 24 hours. To obtain the section using cryostat, bones were frozen in optimum cutting temperature medium (OCT) by placing over dry ice. Femur bones were cut entirely through and made available for analysis. Images were acquired from each bone sections using a confocal microscope connected to Multispectral Imaging System. Later digital images were analyzed and micrometastases were determined.

### Bone marrow cultures

After euthanizing the mice, femur bone was dissected out. Muscles on the bone surface were gently cleaned and then bone marrow was flushed out with RPMI media using 25 gauze needles in a sterilized Eppendorf tube. Cells were dissociated using a syringe and then kept on ice, washed once with RPMI media and then incubated with RPMI media supplemented with 10% FBS, 2 mM glutamine, 100 U/ml penicillin, 100 μg/ml streptomycin, and 50 μg/ml gentamycin. Medium was replaced with fresh RPMI medium twice per week. After 10 days in culture, the presence of ZsGreen fluorescent colonies was determined using an inverted microscope.

### Real-time RT-qPCR

Total RNA from PC3-ML cells was extracted using TRIzol LS Reagent according to the manufacturer’s instructions. RNA quality were assessed by 260/280 ratio by NanoDrop. One microgram of total RNAs was used to generate cDNA using the iScript cDNA synthesis kit (Bio-Rad, Richmond, CA, USA). Real-time PCR was subsequently performed using the iTaq Universal SYBER Green Supermix using the primers listed in Supplemental Table 1 on a CFX96 Real-Time PCR Detection System (Bio-Rad, Richmond, CA, USA). Data were normalized using GAPDH as internal control.

### Immunofluorescence assay

Cells were seeded on glass coverslips coated with Poly-D-lysine (Fisher Scientific). Following treatment, cells were fixed in 4% paraformaldehyde, blocked with 5% normal goat serum containing 0.1% Triton X-100 and then incubated with primary antibody followed by Alexa Fluor 488-conjugated secondary antibody. Images were captured using an Olympus Fluoview 1000 confocal microscope (Bethlehem, PA) equipped with a 60x oil-immersion objective lens (NA, 1.43).

### Statistical analysis

Data analysis was done using the Student’s *t*-test for comparison between two groups (two-tailed). All statistical analyses were done using GraphPad Prism 8 software (GraphPad Software, La Jolla, CA, USA). All values are represented as mean ± standard error of the mean (SEM) of at least three independent experiments. A *p* value < 0.05 was considered statistically significant (* *p* < 0.05, ** *p* < 0.01, *** *p* < 0.001, **** *p* < 0.0001, ns: not significant).

## Results

### The expression and activity of PKD2 and PKD3 are required for the viability, migration, invasion of prostate cancer cells

PKD has emerged as a promising therapeutic target for cancer including prostate cancer. However, its precise role in prostate cancer metastasis, particularly bone metastasis, and the mechanisms through which it regulates this process remain largely unknown. We seek to define the role of PKD tumor metastasis to the bone by using a PC3-ML bone metastasis model. PC3-ML, a subline of PC3, displays 80% metastatic efficiency to the lumbar vertebrae and produces bone metastasis when injected in mice (21). First, we examined the expression of PKD isoforms in PC3-ML, as compared to a normal prostate epithelial cell line RWPE-1 and an androgen-dependent prostate cancer cell lines LNCaP by Western blotting (Fig. 1A). RWPE-1 and LNCaP cells expressed all three PKD isoforms, while PC3-ML only expressed PKD2 and PKD3. Next, we examined the functional input of PKD2/3 to PC3-ML proliferation and migration/invasion. PKD2 and PKD3 were silenced by two separate siRNAs that targeted different regions of the PKD2 or 3 genes, and their activities were inhibited by a pan-PKD inhibitor CRT0066101 (CRT101). The cells were then examined for long-term colony forming ability. As shown in Fig. 1B, our results showed that silencing of PKD2 or PKD3 dramatically decreased the number of colonies formed as compared to the control, and inhibition of PKD by CRT101 had similar effect. Quantitative analysis further confirmed these findings (Fig. 1B, *left*). Accordingly, overexpression of a catalytic active PKD2 mutant (PKD2-CA) promoted the formation of colonies by 2-fold as compared to the empty vector control (Fig. 1C). Taken together, these data indicated that PKD2/3 played an essential role in colonigenic potential of PC3-ML cells.

**Figure 1.**
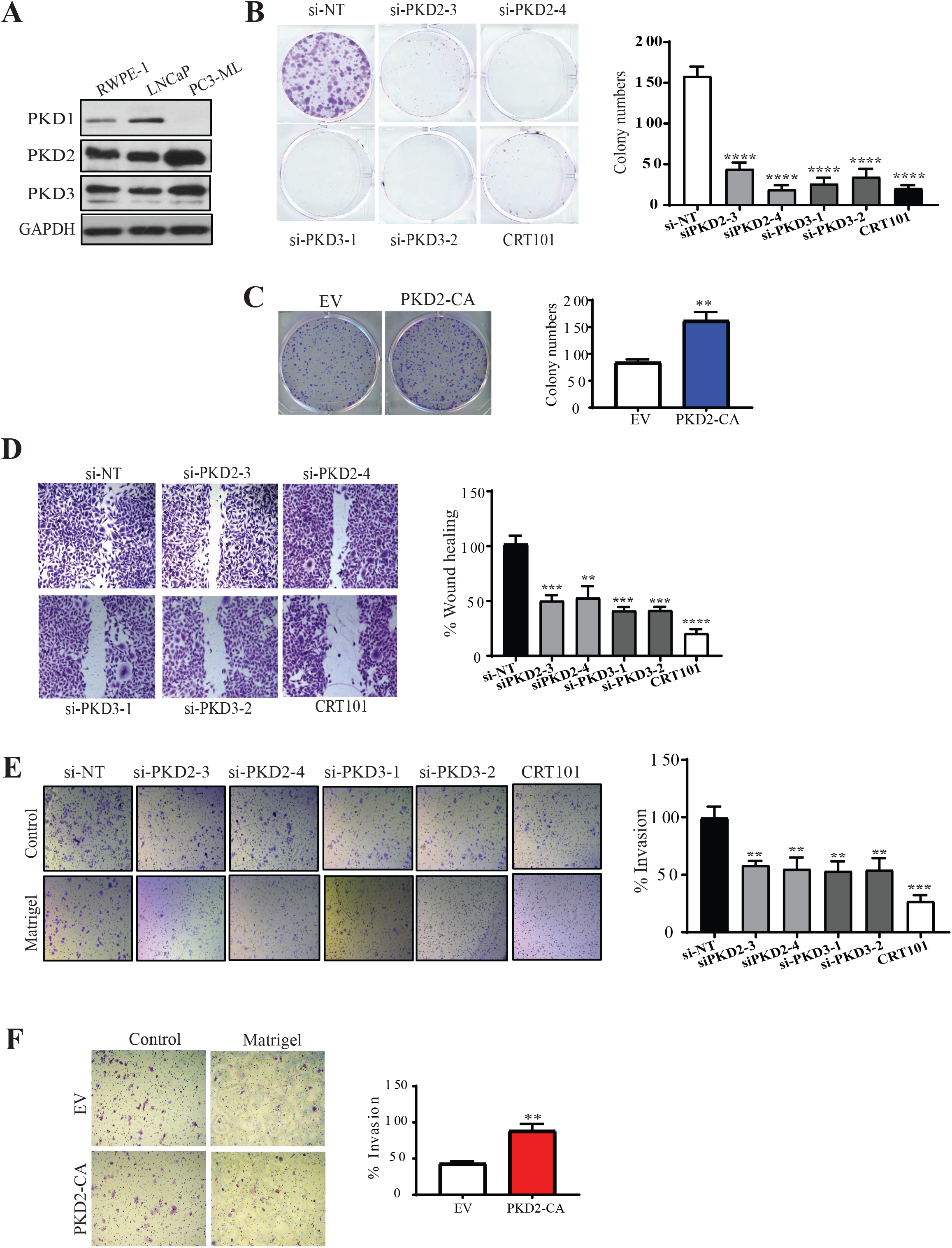
Depletion of PKD2/3 or inhibition of PKD activity suppressed the proliferation, migration, and invasion of PC3-ML prostate cancer cells. (**A**) Expression of PKD1, 2, 3 was analyzed in RWPE-1, LNCaP, and PC3-Ml cells by Western blotting. (**B-C**) Depletion or inhibition of PKD blocked colonigenic ability of PC3-ML cells. PC3-ML (1000 cells per well) were transfected with PKD2 or PKD3 siRNAs, or exposed to CRT101 (5μM) (B), or transfected with an empty vector (EV) or a catalytically active PKD2 (PKD2-CA) plasmid (C). Two days later, the medium was replaced with fresh medium. After 9 days, the cells were stained with crystal violet and were counted for colony formation. *right panel*, colony numbers in 5 random fields. (**D**) Depletion or inhibition of PKD inhibited PC3-ML prostate tumor cell migration and invasion. Cells were transfected with PKD2 or PKD3 siRNAs for 48 h or treated with CRT0066101 (5μM) for 24 h. Cells were stained with crystal violet (*left panel*) and wound closure was measured by Image J (*right panel*). (**E**) Depletion or inhibition of PKD blocked PC3-ML prostate tumor cell invasion. PC3-ML cells were transfected with PKD2 or PKD3 siRNAs for 48 h, then plated into transwell inserts coated with or without Matrigel. Additionally, cells were treated with the PKD inhibitor CRT0066101 (5μM) in transwell inserts. After 22 h, noninvasive cells were removed and invasive cells were stained, photographed, and quantified counts in 5 random fields. *Right panels*, percentage invasion was calculated as the percent of the cells invaded through Matrigel inserts vs. the total cells migrated through the control inserts. (**F**) Matrigel invasion Assay was performed on cells transfected with a control (EV) or a PKD2-CA plasmid. Cells invaded through Matrigel were imaged (*left panel*) and quantified (*right panel*). Data shown in the bar graphs are the average of three independent experiments with error bars representing SEM (* p<0.05, ** p < 0.01, *** p < 0.001, **** p < 0.0001).

We then determined whether silencing of PKDs in PC3-ML cells inhibited the migratory and/or invasive properties of the tumor cells. PC3-ML cells were transfected with PKD2/3 siRNAs or treated with CRT0066101, followed by wound healing assay. We found that silencing of PKDs resulted in 22% to 35% inhibition of cell migration as compared to control (Fig. 1D). Incubation of cells with CRT101 for 24 h resulted in 87% inhibition of wound closure as compared to the control. This data indicated that PKD2 and PKD3 along with its inhibitor potently decreased the migration of prostate cancer cells.

We further determined whether depletion of PKDs also modulated the invasion of prostate cancer cells. We found that silencing of PKD2 and PKD3 in PC3-ML for 22 h resulted in about 40% inhibition of cell invasion compared with control (Fig. 1E). Also, inhibition of PKD by CRT0066101 resulted in over 60% decreased invasion of PC3-ML cells. Accordingly, overexpression of a catalytic active PKD2-CA doubled the number of cells invaded through the Matrigel (Fig. 1F). Thus, our data clearly showed that PKD2 and 3 were required for prostate cancer cell invasion *in vitro*.

### PKD2 and 3 modulated the expression of bone metastatic genes in prostate cancer cells

A number of metastatic genes including osteopontin (OPN), osteocalcin (OCN), alkaline phosphatase (ALP), integrin 1β, talin1, focal adhesion kinase (FAK), matrix metaloprotienase-9, interleukin 1α and interleukin 1β are known to contribute to bone metastasis of cancer cells. These molecules regulate the process of bone metastasis and the phenomenon of tumor dormancy in the bone marrow as well as create metastatic niches (23,24). We evaluated the impact of silencing PKD2/3 on the expression of a panel of these metastasis-related genes in prostate cancer cells. We found that several bone metastatic genes including FAK, OCN and OPN were decreased as a result of PKD2/3 silencing (Fig 2A). As negative controls, collagenase I did not show any significant change in expression either by silencing of PKD2 and PKD3 or by CRT0066101 treatment (data not shown).

**Figure 2.**
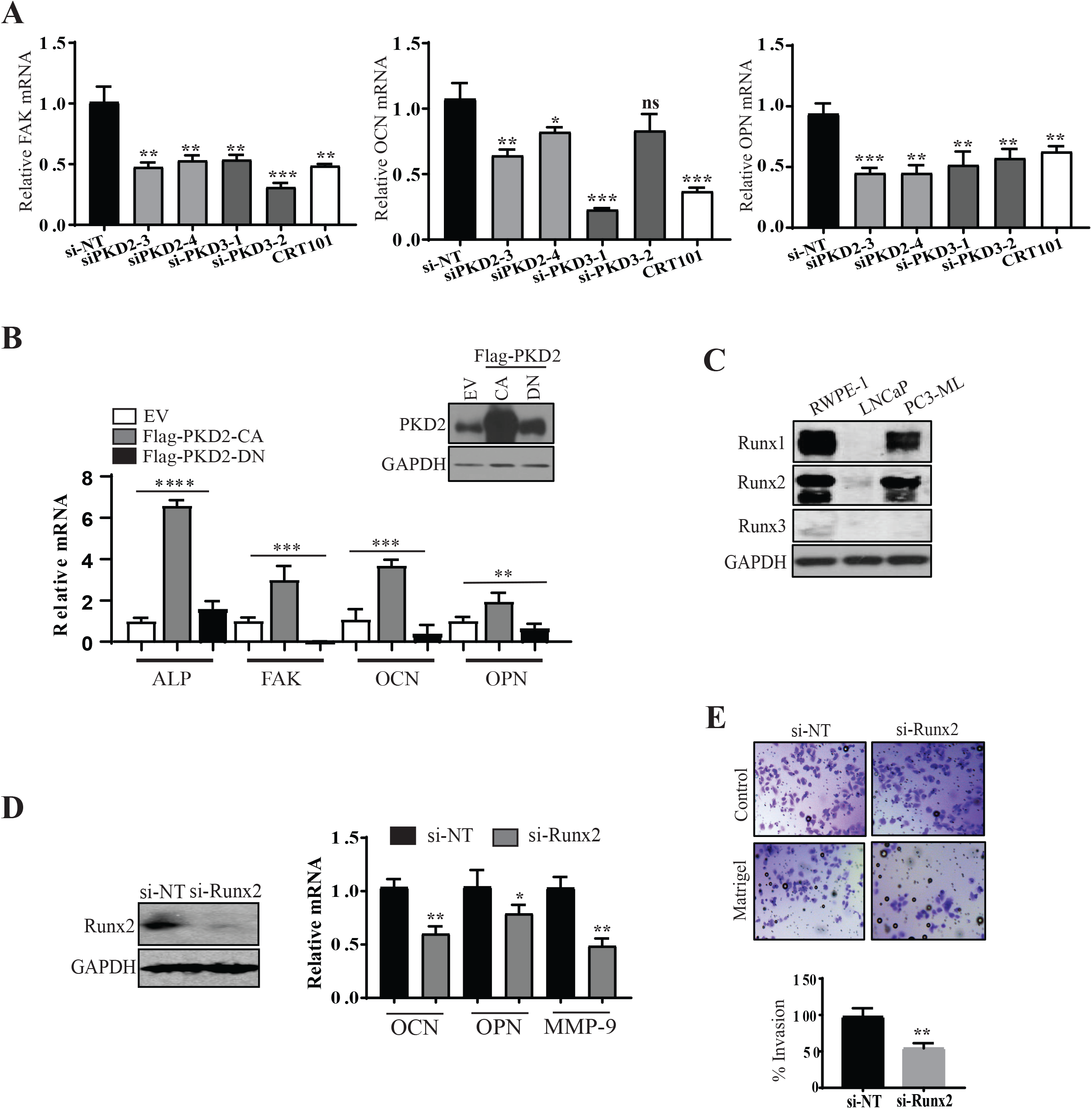
Depletion or inhibition of PKD blocked the expression of Runx2-regualted bone metastatic genes in prostate cancer cells. (**A**) PC3-ML cells were transfected with PKD2 or PKD3 siRNA or treated with CRT101 (5μM) for 24 h. RT-qPCR was performed for the genes as shown. Transcript levels relative to the GAPDH control were determined in PC3-ML cells. (**B**) PC3-ML cells were transfected with empty vector (EV), Flag-PKD2-CA and Flag-PKD2-DN. Transcript levels of ALP, FAK, OCN, and OPN were determined by real time RT-qPCR. *inset*, confirmation of the overexpressed PKD2-CA and -DN in PC3-ML by western blotting. (**C**) Runx expression in prostate cancer cells. Whole cell extracts of RWPE-1, LNCaP and PC3-ML cells were subjected for western blotting for Runx1-3 expression. (**D**) Runx2 regulated the expression of OCN, OPN and MMP-9. PC3-ML were transfected with a control siRNA (si-NT) or a Runx2 siRNA (si-Runx2), followed by real time RT-qPCR for the indicated genes. *left*, western blot confirmed the knockdown of Runx2. (**E**) Matrigel invasion assay were performed onPC3-ML cells transfected with si-NT and si-Runx2. Percent invasion was calculated and shown in the bar graph. For all the qPCR experiments, each experiment was repeated with duplicates at least 2–3 times and data represent the Mean ± SEM from all independent experiments.

The impact of PKD2 on the expression of bone metastatic genes was further examined by overexpressing the catalytic active PKD2-CA and a dominant negative PKD2-DN in PC3-ML cells. As shown in Fig. 2B, overexpression of PKD2-CA increased the expression of ALP, FAK, OCN, and OPN, while PKD2-DN blocked or did not affect the expression of these genes. The overexpression of PKD2-CA and -DN proteins were confirmed by western blotting (Fig. 2B, upper panel). The regulation of bone metastasis genes by PKD2 and 3 implied a potential role of PKD in prostate cancer bone metastasis.

### PKDs regulated bone metastatic gene expression through Runx2

Runx2 knockdown is known to decrease cell migratory and invasive ability of cancer cells (25). Therefore, we investigated whether silencing of Runx2 inhibits invasion of prostate cancer cells. First, we determined the expression level of Runx1, Runx2 and Runx3 in PC3-ML cells and compared with other cell lines including prostate epithelial RWPE-1 and androgen-dependent LNCaP prostate cancer cells. We found that Runx1 and Runx2 are highly expressed in RWPE-1 and PC3-ML cell while Runx3 was undetectable in all three cell lines (Fig 2C). We also confirmed at mRNA level by qPCR that Runx1 and Runx2 mRNA level is higher in PC3-ML cells compared to LNCaP cells (Data not shown). Runx2 is a key regulator of bone metastasis in multiple cancers including prostate (20), therefore we sought to elucidate the role of Runx2 in PKD-regulated tumor metastasis in prostate cancer cell. We first examined whether Runx2 was directly involve in metastatic gene expression in prostate cancer cells. As shown in Fig. 2D, silencing Runx2 resulted in the downregulation of the same bone metastasis-associated genes, MMP-9, OCN, and OPN, that were regulated by PKD2/3. Functionally, knockdown of Runx2 resulted in around 20% inhibition of cell invasion compared with control (Fig. 2E). Thus, Runx2 contributed to the invasive properties of prostate cancer cells.

### Depletion or inactivation of PKD accelerated Runx2 degradation

To gain insight into the regulation of Runx2 by PKD in prostate cancer cells, we examined the effects of altering PKD expression or activity on Runx2 mRNA and protein expressions. As shown in Fig. 3A, Knockdown of PKD2 by siRNAs or inactivating PKD by CRT0066101 resulted in a small decrease in Runx2 mRNA levels, while knockdown of PKD3 did not significantly affect Runx2 transcription (Fig. 3A). Meanwhile, depletion or inactivation of PKD had no effect on Runx1 expression (Fig. S1A). The siRNAs were effective at knocking down PKD2 and PKD3 (Fig. S1B-C). These data indicate that PKD2/3 had a marginal effect in Runx2 transcription in prostate cancer cells. Next, we examined the expression and localization of Runx2 by immunoblotting and immunofluorescence (IF) staining before or after the knockdown of PKD2 or PKD3 and inhibition of PKD activity by CRT101. As shown in the Fig. 3B, knocking down or inactivating PKD significantly downregulated Runx2 at protein level in PC3-ML cells. Accordingly, the overall intensity of Runx2 staining was significantly diminished upon depletion of PKD2/3 or inactivation of PKD, while its nuclear localization was not altered (Fig. 3C). These data suggest that PKD regulated Runx2 protein expression mainly at post-transcriptional level in prostate cancer cells. To further this analysis, we examined if PKD may regulate the stability of Runx2 protein. The cells were treated with either DMSO or CRT101 over different times in the presence of cycloheximide, an inhibitor of protein synthesis, the levels of Runx2 protein were examined by Western blotting (Fig. 3D). Our data showed that inhibition of PKD by CRT101 accelerated the degradation of Runx2. The protein half-life of Runx2 in presence of CRT101 was reduced to 3.2 h from 17.6 h (DMSO) (Fig. 3D, *lower panel*). Taken together, PKD positively regulates Runx2 at post-translational levels by stabilizing Runx2 protein.

**Figure 3.**
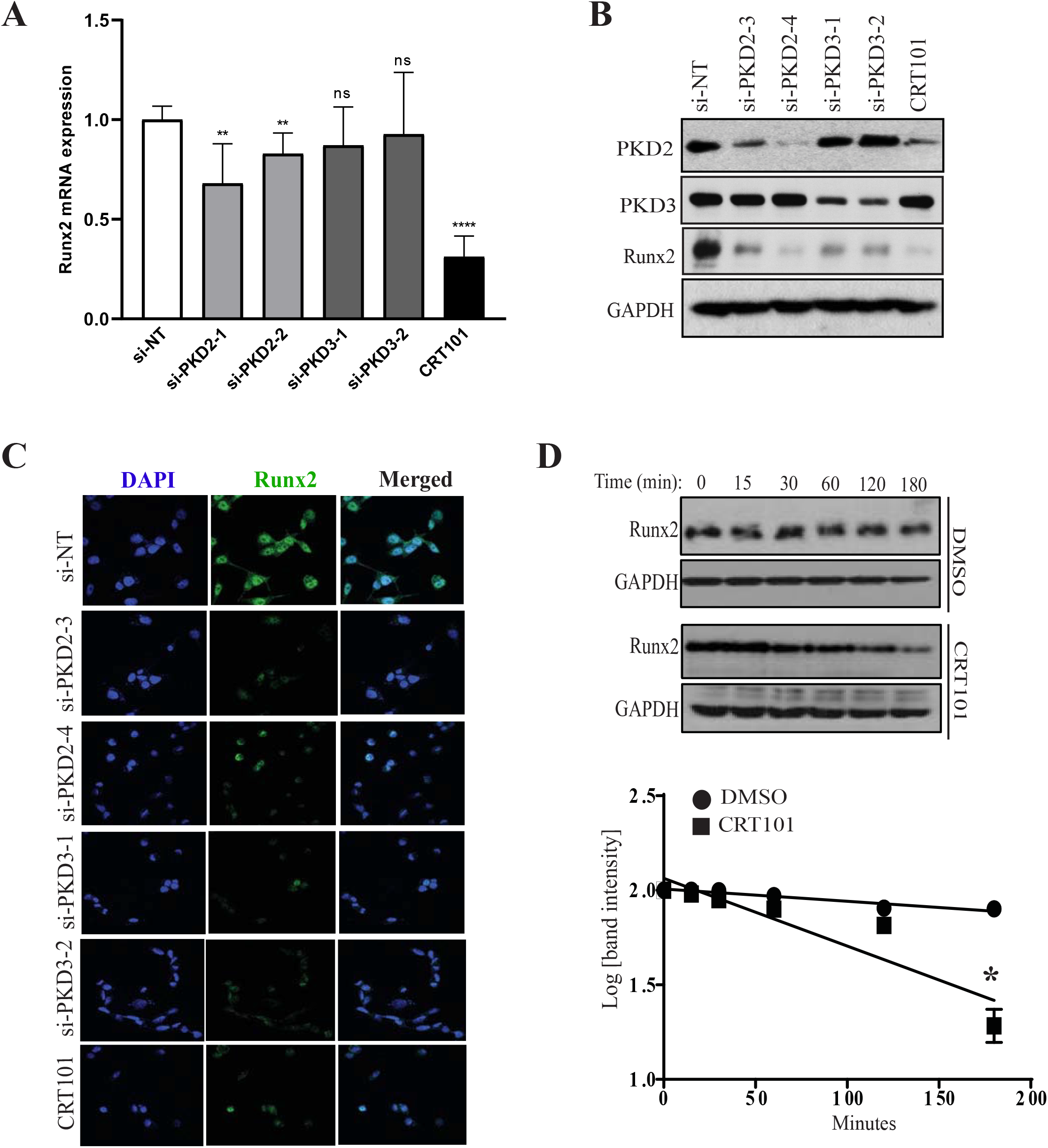
Inactivation or knockdown of PKD2 causes downregulation of Runx2 protein in prostate cancer cells. (**A-B**) Transcript and protein levels of Runx1, 2 and PKD2, 3 were analyzed by real time RT-qPCR (A) or western blotting (B) in PC3-ML cells transfected with PKD2 or PKD3 siRNA or treated with CRT101 (5μM). (**C**) Immunofluorescence staining was performed on tumor cells transfected with PKD2 and PKD3 siRNAs or treated with CRT101 using Runx2 antibody. The nuclei were counterstained with DAPI. Representative images from three independent experiments are shown. (**D**) PC3-ML Cells were treated with cycloheximide with or without CRT101 for indicated times and cell lysates were immunoblotted for Runx2. GAPDH served as loading control. *bottom*, half-life of Runx2 protein was measured from the blots above. Data are the Mean ± SEM of three independent experiments (* p<0.5, ** p < 0.01, *** p < 0.001, ns: not significant).

### PKD modulated Runx2 activity through Ras/ERK1/2 MAPK signaling pathway

Emerging evidence suggests that PKDs modulate a plethora of cellular functions involved in tumorigenesis via Ras/Raf/MEK/ERK1/2 MAPK signaling cascade (26). To gain mechanistic insights into PKD-regulated Runx2 activity in bone metastasis of prostate cancer, we determined whether PKDs modulated Runx2 and downstream target genes responsible for prostate cancer bone metastasis via MEK/ERK1/2 axis. PC3-ML cells were transfected with empty vector (EV) or a plasmid carrying constitutively active PKD2 (Flag-PKD2^CA^) and activation of ERK1/2 and expression of Runx2 were determined by Western blotting (Fig. 4A). Indeed, overexpression of active PKD2 increased both phosphor-ERK1/2 and Runx2 levels and this effect was dampened when PC3-ML cells were treated with U0126, a highly selective inhibitor of MAPK/ERK kinases (Fig. 4B). In line of these observations, we found that treatment of PC3-ML cells with U0126 significantly downregulated the mRNA levels of Runx2 and several genes responsible for bone metastasis including FAK, OCN and OPN (Fig. 4C). Moreover, overexpression of PKD2 in U0126-treated PC3-ML cells rescued the dampened expressions of Runx2, ALP, FAK, OCN and OPN caused by U0126 as analyzed by both qRT-PCR and Western blotting (Fig. 4D). Taken together, these results suggested that PKD modulates Runx2 activity through MEK/ERK1/2 MAPK signaling pathway.

**Figure 4.**
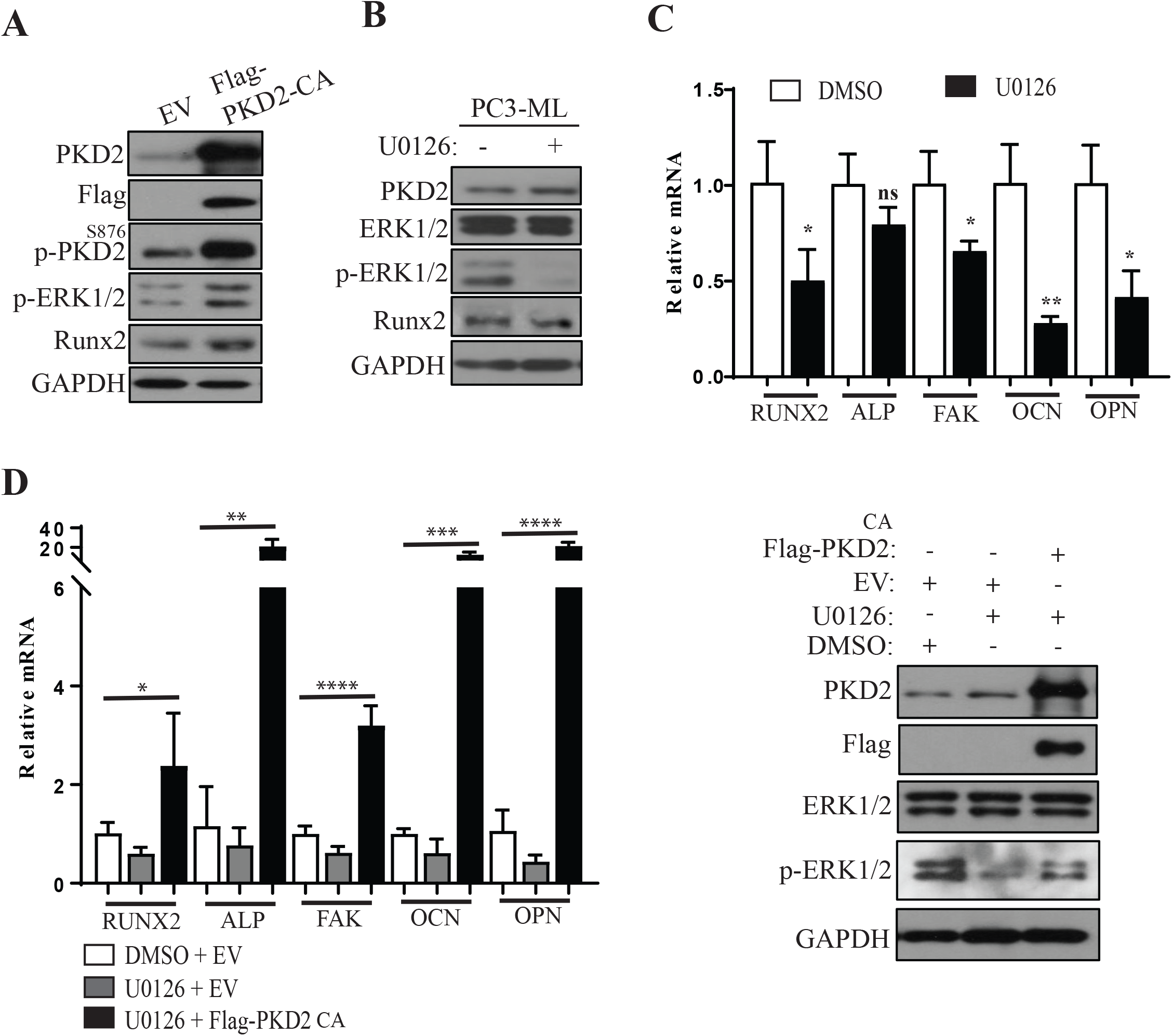
PKD regulated bone metastatic gene expression through MEK/ERK. (**A**) Overexpression of PKD2-CA increased ERK activity. PC3-ML cells were transfected with empty vector (EV) or Flag-PKD2-CA, followed by western blotting for indicated genes. (**B**) U0126 blocked ERK1/2 phosphorylation in PC3-ML cells. (**D-C**) PC3-ML cells were treated with the MEK inhibitor U0126. Transcript levels of Runx2, ALP, FAK, OCN, and OPN were determined by real time RT-qPCR. Data are the Mean ± SEM of three independent experiments (* p<0.5, ** p < 0.01, *** p < 0.001, ns: not significant).

### Inhibition of PKD blocked accumulation of PC3-ML cells in the mouse bone marrow and reduced tumor metastasis of prostate cancer cells to bone in mice

PKD has been implicated in a variety of cancer-associated biological processes including cancer cell growth, apoptosis, motility, and angiogenesis (4). However, its role in bone metastasis *in vivo* in the context of prostate cancer has not been shown. Here, using a mouse model of bone metastasis generated by cardiac injection of tumor cells, we determined the potential role of PKD in bone metastasis by targeted inhibition of PKD using CRT101. This is a valuable model for bone metastasis research as well as standard and time-honored technique to investigate human cancer bone metastasis *in vivo*. Use of fluorescent cells is required to detect the circulation and homing of cells in the bone. Therefore, we used fluorescent PC3-ML ZSGreen cells and inoculated in the left cardiac ventricle of athymic nude mice, as described elsewhere (27). PC3-ML ZSGreen cell lines were produced by stabliy expressing ZsGreen in the PC3-ML cells(27). After injection of cells, animals were randomly divided into two treatment groups: vehicle and CRT101. Treatment of vehicle and PKD inhibitor CRT101 (80 mg/kg) were given orally daily for 4 weeks (Fig 5A). At the end of experiment, animals were sacrificed and femur bones from each mouse were dissected out. One of the femur bones from each mouse used for decalcification followed by cryosectioning and bone marrow of other femur bone was flushed out and cultured in RPMI media. The body weight of animals that was recorded twice a week showed no significant change during the experiment (Fig 5B).

**Figure 5.**
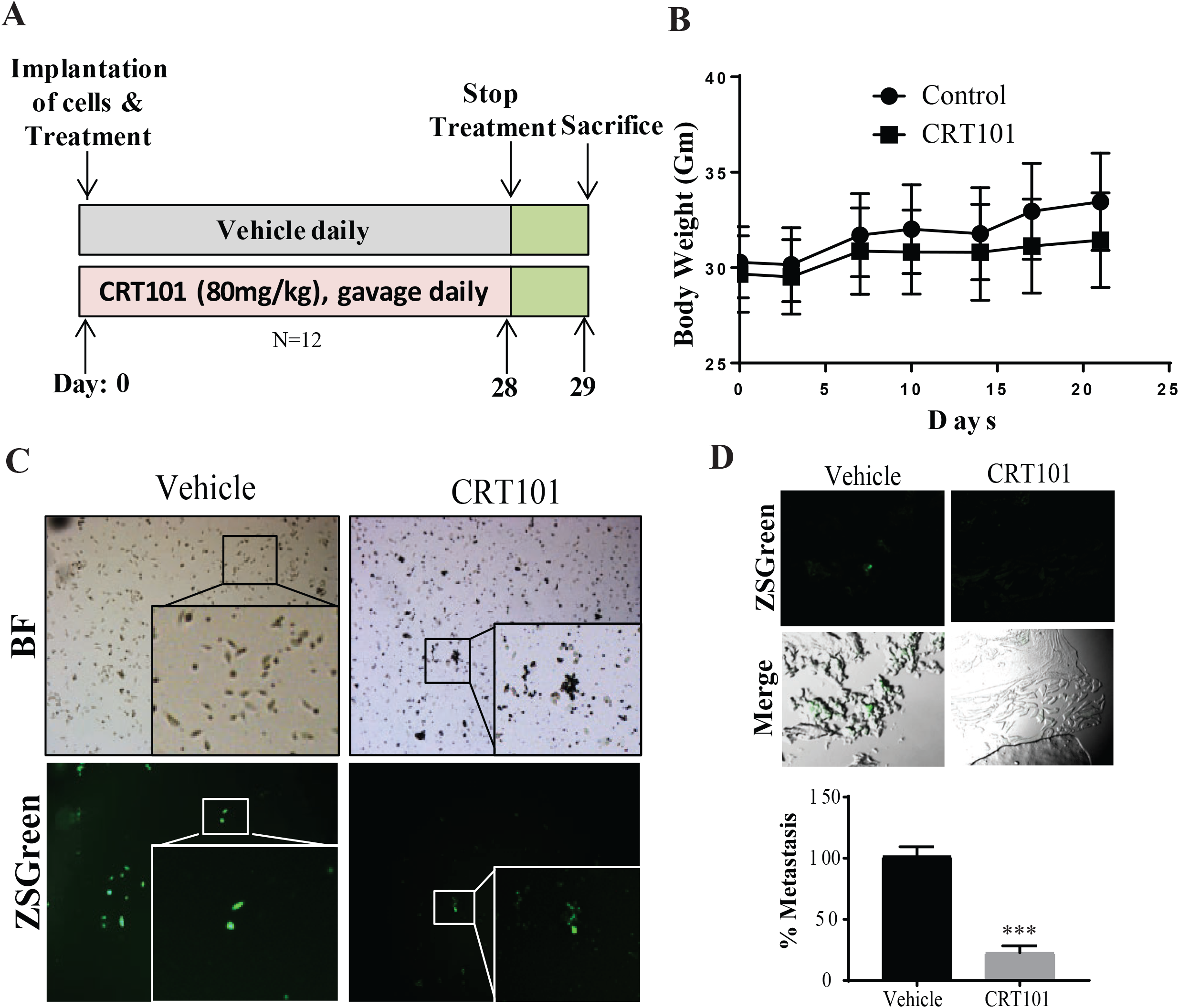
PKD inhibitor CRT0066101 impairs metastatic dissemination of PC3-ML cells to the athymic mouse bone. (**A**) Schematic representation for the treatment of mice. Male athymic nude mice (n=12) were inoculated by intracardiac injection with the ZsGreen PC3-ML cell lines. Animals were treated with CRT101 (80 mg/kg, gavage daily) and vehicle (5% dextrose). (**B**) Body weight was recorded twice a week. (**C**) After 28 days, skeletal micrometastasis was identified by confocal microscopy. A representative picture showing ZsGreen PC3-ML cell micrometastases (left panel). Percent decrease in skeletal micrometastasis on CRT101 treated animals after 28 days of injection. *Significant (P<0.05) over controls (right panel). (**D**) Twenty eight days after intracardiac injection, cells from bone marrow were flushed and cultured for 10 days. Representative micrographs are shown.

First, we determined whether CRT101 affected the migration and accumulation of PC3-ML cells in bone marrows. The presence of fluorescent ZsGreen labeled cells in bone marrows was analyzed in mice treated with or without CRT101. At the end of experiment, bone marrows cells were flushed out from femurs and cultured *in vitro*. After ten days of culture we found a significant number of ZsGreen labeled cells in bone marrows from mice treated with vehicle as compared to ZsGreen fluorescent cells detected in bone marrow of CRT101-treated mice (Fig. 5C).

It has been shown that upon direct inoculation of prostate cancer cells in the blood circulation of immunodeficient mice, these cells generate metastatic foci primarily in femur and tibia, jaws and ribs (21). In view of this, we inoculated PC3-ML cells in the left cardiac ventricle of athymic nude mice as described above. After completion of treatment period of 28 days, mice were sacrificed and their femur harvested, fixed, and analyzed for the presence of micrometastatic foci. As shown in figure 5D, we found microscopic tumors in the femur upon intracardiac inoculation of PC3-ML ZsGreen cells. Quantification analysis revealed that treatment of CRT101 in PC3-ML ZsGreen inoculated mouse severely impaired the formation of micrometastasis in the bone (Fig. 5D, *lower graph*). Thus, this result indicated that inhibition of PKD impairs accumulation of prostate cancer cells.

## Discussion

Metastasis is the spread of cancer cells from the primary tumor, which initiate new tumors to surrounding tissues and to distant organs. Till date a number of signaling molecules have been discovered that contributes to the metastasis of different cancer cells. However, the mechanisms that lead to prostate cancer bone metastasis are not completely known. Although PKD has known to play important roles in cell proliferation, survival, migration, gene regulation, protein trafficking, and immune response, its role in prostate bone metastasis has not been investigated. In this study we identified PKD as a potential mediator of prostate cancer bone metastasis. We found that depletion of PKD2/3 or inhibition of PKD blocked tumor proliferation, migration and invasion at cellular levels and inhibition of PKD reduced the formation of tumor micrometastases in a mouse bone metastasis model.

Several lines of evidence support that PKDs contributes to the proliferation of multiple cancer cell types including prostate cancer (16,28,29). In the present study, to investigate the effect of PKD2/3 on these properties, we inhibited expression of PKD2/3 by using respective siRNAs and PKD inhibitor CRT0066101 in PC3-ML cells, which naturally expresses PKD2 and PKD3. We found that depletion of PKD2/3 resulted in suppression of cell growth and proliferation, which agrees with our previous findings (19). Furthermore, this result was further substantiated by pharmacological inhibition of PKD activity with a pan-PKD inhibitor CRT101, which reduced colony formation. Our results indicate that PKD2/3 plays an important role in prostate cancer cell proliferation.

As PKD2 and PKD3 coordinate to promote prostate cancer cell invasion (17), they may directly involve in invasion and migration of cancer cells. In this context, we silenced the PKDs and analyzed the migration of PC3-ML prostate cancer cells by wound healing and invasion assay.

We found that depletion of PKD2 and PKD3 in prostate cancer PC3-ML cells reduced the migration and invasion of cells. Although PC3-ML does not express PKD1, overexpression of PKD1 in prostate cancer LNCaP cells increases aggregation, but reduces motility and invasiveness through interaction with and phosphorylation of E-cadherin (30). Other study showed that PKD2 silencing inhibits migration of doxorubicin resistant MCF7 cells without affecting chemoresistance (31). Deng group also reported PKD2/PKD3 coordinate to promote prostate cancer cell invasion through p65 NF-κB and expression and activation of uPA (17). In this study, we found that depletion of PKD results in decrease in another transcription factor Runx2. We specifically emphasized on Runx2 because it involves in prostate cancer bone metastasis (20). Runx2 is a member of mammalian Runt homology domain transcription factors. Runx proteins are involved in cell lineage determination during development and various forms of cancer (32). Runx2 is critical for regulation of genes that support bone formation and is abnormally expressed in tumors that metastasize to the bone (33). It activates several metastatic genes including MMPs, VEGF, osteopontin, RANKL in metastatic cancer cells (34). We also found that depletion of Runx2 blocked migration/invasion of prostate cancer cells and suppressed expression of various bone metastasis-associated genes in PC3-ML cells. Our data suggest that Runx2 can act either independently or downstream of PKD2/3 to inhibit metastasis of prostate cancer cells. We also found that the PKD inhibitor CRT101 suppressed invasion and metastasis markers in prostate cancer cells. This finding has been supported by several reports. Borges et al (35) showed that treatment with CRT101 inhibited growth of primary breast tumors and metastasis in both *in vivo* and *in vitro*. CRT101 induced apoptosis in pancreatic cancer cells mediated through suppression of NF-κB pathway. It also inhibits growth of pancreatic tumor in xenograft model, correlating to changes of cyclin D1, survivin, and cIAP-1 (36). It has also shown to inhibit migration and invasion of U87MG cells by inhibiting p44/42 MAPK and to a smaller extent p54/46 JNK and p38 MAPK activation (37). In this study, we found that CRT101 suppressed invasion and migration of prostate cancer cells by blocking Runx2 expression at both transcriptional (minor effect) and post-transcriptional (major effect) levels. Importantly, we also found that this inhibitor CRT101 suppressed the bone metastasis of prostate cancer cells in a mouse metastasis model using intracardiac injection. These results indicate that inhibition of PKD either by genetic depletion or pharmacological inhibition suppressed the bone metastasis of prostate cancer cells.

Conclusively, our study identified that PKD2/3 has important role in prostate cancer metastasis through the transcription factor Runx2. For the first time, we also found that a PKD inhibitor CRT101 inhibits the migration and accumulation of prostate cancer cells to the bone in a mouse metastasis model. These findings may have significant therapeutic implications against bone metastatic diseases, as CRT101 has shown promising anticancer activities against various cancers (35,36,38).

## Acknowledgment

We would like to thank Dr. Alvaro Gutierrez-Uzquiza in the Department of Systems Pharmacology and Translational Therapeutics, University of Pennsylvania, Philadelphia for invaluable discussions and guidance on intracardiac injection of prostate cancer cells in animals. We are also grateful for the technical assistance from Dr. Xuejing Zhang on QPCR analysis.

## References

1. Patrikidou A, Loriot Y, Eymard JC, Albiges L, Massard C, Ileana E, et al. Who dies from prostate cancer? Prostate cancer and prostatic diseases 2014;17(4):348–52 doi 10.1038/pcan.2014.35.

2. Bubendorf L, Schopfer A, Wagner U, Sauter G, Moch H, Willi N, et al. Metastatic patterns of prostate cancer: an autopsy study of 1,589 patients. Human pathology 2000;31(5):578–83.

3. Tsuzuki S, Park SH, Eber MR, Peters CM, Shiozawa Y. Skeletal complications in cancer patients with bone metastases. International journal of urology : official journal of the Japanese Urological Association 2016;23(10):825–32 doi 10.1111/iju.13170.

4. Roy A, Ye J, Deng F, Wang QJ. Protein kinase D signaling in cancer: A friend or foe? Biochimica et biophysica acta 2017;1868(1):283–94 doi 10.1016/j.bbcan.2017.05.008.

5. Sharlow ER, Giridhar KV, Lavalle CR, Chen J, Leimgruber S, Barrett R, et al. Potent and Selective Disruption of Protein Kinase D Functionality by a Benzoxoloazepinolone. J Biol Chem 2008;283(48):33516–26 doi 10.1074/jbc.M805358200.

6. Guo JX, Clausen DM, Beumer JH, Parise RA, Egorin MJ, Bravo-Altamirano K, et al. In vitro cytotoxicity, pharmacokinetics, tissue distribution, and metabolism of small-molecule protein kinase D inhibitors, kb-NB142-70 and kb-NB165-09, in mice bearing human cancer xenografts. Cancer Chemoth Pharm 2013;71(2):331–44 doi 10.1007/s00280-012-2010-z.

7. Tandon M, Wang LR, Xu Q, Xie XQ, Wipf P, Wang QJ. A Targeted Library Screen Reveals a New Inhibitor Scaffold for Protein Kinase D. PloS one 2012;7(9) doi ARTN e44653 10.1371/journal.pone.0044653.

8. Tandon M, Johnson J, Li ZH, Xu SP, Wipf P, Wang QJ. New Pyrazolopyrimidine Inhibitors of Protein Kinase D as Potent Anticancer Agents for Prostate Cancer Cells. PloS one 2013;8(9) doi ARTN e75601 10.1371/journal.pone.0075601.

9. Tandon M, Salamoun JM, Carder EJ, Farber E, Xu SP, Deng F, et al. SD-208, a Novel Protein Kinase D Inhibitor, Blocks Prostate Cancer Cell Proliferation and Tumor Growth In Vivo by Inducing G2/M Cell Cycle Arrest. PloS one 2015;10(3) doi ARTN e0119346 10.1371/journal.pone.0119346.

10. Li QQ, Hsu I, Sanford T, Railkar R, Balaji N, Sourbier C, et al. Protein kinase D inhibitor CRT0066101 suppresses bladder cancer growth in vitro and xenografts via blockade of the cell cycle at G2/M. Cell Mol Life Sci 2018;75(5):939–63 doi 10.1007/s00018-017-2681-z.

11. Borges S, Perez EA, Thompson EA, Radisky DC, Geiger XJ, Storz P. Effective Targeting of Estrogen Receptor Negative Breast Cancers with the Protein Kinase D inhibitor CRT0066101. Mol Cancer Ther 2015 doi 10.1158/1535-7163.MCT-14-0945.

12. Wei N, Chu E, Wipf P, Schmitz JC. Protein kinase d as a potential chemotherapeutic target for colorectal cancer. Mol Cancer Ther 2014;13(5):1130–41 doi 10.1158/1535-7163.MCT-13-0880.

13. Harikumar KB, Kunnumakkara AB, Ochi N, Tong Z, Deorukhkar A, Sung B, et al. A novel small-molecule inhibitor of protein kinase D blocks pancreatic cancer growth in vitro and in vivo. Mol Cancer Ther 2010;9(5):1136-46 doi 1535-7163.MCT-09-1145 [pii] 10.1158/1535-7163.MCT-09-1145.

14. Zhang LY, Zhao ZL, Xu SP, Tandon M, LaValle CR, Deng F, et al. Androgen suppresses protein kinase D1 expression through fibroblast growth factor receptor substrate 2 in prostate cancer cells. Oncotarget 2017;8(8):12800–11 doi 10.18632/oncotarget.14536.

15. Biswas MH, D. C, Zhang C, Straubhaar J, Languino LR, Balaji KC. Protein kinase D1 inhibits cell proliferation through matrix metalloproteinase-2 and matrix metalloproteinase-9 secretion in prostate cancer. Cancer Res 2010;70(5):2095-104 doi 0008-5472.CAN-09-4155 [pii] 10.1158/0008-5472.CAN-09-4155.

16. Chen J, Deng F, Singh SV, Wang QJ. Protein kinase D3 (PKD3) contributes to prostate cancer cell growth and survival through a PKCepsilon/PKD3 pathway downstream of Akt and ERK 1/2. Cancer Res 2008;68(10):3844–53 doi 10.1158/0008-5472.CAN-07-5156.

17. Zou ZP, Zeng FY, Xu WF, Wang CX, Ke ZY, Wang QJ, et al. PKD2 and PKD3 promote prostate cancer cell invasion by modulating NF-kappa B-and HDAC1-mediated expression and activation of uPA. J Cell Sci 2012;125(20):4800–11 doi 10.1242/jcs.106542.

18. Zou Z, Zeng F, Xu W, Wang C, Ke Z, Wang QJ, et al. PKD2 and PKD3 promote prostate cancer cell invasion by modulating NF-κB-and HDAC1-mediated expression and activation of uPA. J Cell Sci 2012;125(Pt 20):4800–11 doi 10.1242/jcs.106542.

19. LaValle CR, Zhang L, Xu S, Eiseman JL, Wang QJ. Inducible silencing of protein kinase D3 inhibits secretion of tumor-promoting factors in prostate cancer. Mol Cancer Ther 2012;11(7):1389–99 doi 10.1158/1535-7163.MCT-11-0887.

20. Akech J, Wixted JJ, Bedard K, van der Deen M, Hussain S, Guise TA, et al. Runx2 association with progression of prostate cancer in patients: mechanisms mediating bone osteolysis and osteoblastic metastatic lesions. Oncogene 2010;29(6):811–21 doi 10.1038/onc.2009.389.

21. Wang M, Stearns ME. Isolation and characterization of PC-3 human prostatic tumor sublines which preferentially metastasize to select organs in S.C.I.D. mice. Differentiation; research in biological diversity 1991;48(2):115–25.

22. Chen J, Deng F, Singh SV, Wang QJ. Protein kinase D3 (PKD3) contributes to prostate cancer cell growth and survival through a PKC epsilon/PKD3 pathway downstream of Akt and ERK 1/2. Cancer Research 2008;68(10):3844–53 doi 10.1158/0008-5472.can-07-5156.

23. Weidle UH, Birzele F, Kollmorgen G, Ruger R. Molecular Mechanisms of Bone Metastasis. Cancer Genom Proteom 2016;13(1):1–12.

24. Jin JK, Tien PC, Cheng CJ, Song JH, Huang C, Lin SH, et al. Talin1 phosphorylation activates beta 1 integrins: a novel mechanism to promote prostate cancer bone metastasis. Oncogene 2015;34(14):1811–21 doi 10.1038/onc.2014.116.

25. Boregowda RK, Olabisi OO, Abushahba W, Jeong BS, Haenssen KK, Chen WJ, et al. RUNX2 is overexpressed in melanoma cells and mediates their migration and invasion. Cancer Lett 2014;348(1-2):61–70 doi 10.1016/j.canlet.2014.03.011.

26. Zhang X, Connelly J, Chao Y, Wang QJ. Multifaceted Functions of Protein Kinase D in Pathological Processes and Human Diseases. Biomolecules 2021;11(3) doi 10.3390/biom11030483.

27. Russell MR, Jamieson WL, Dolloff NG, Fatatis A. The alpha-receptor for platelet-derived growth factor as a target for antibody-mediated inhibition of skeletal metastases from prostate cancer cells. Oncogene 2009;28(3):412–21 doi 10.1038/onc.2008.390.

28. Huck B, Duss S, Hausser A, Olayioye MA. Elevated Protein Kinase D3 (PKD3) Expression Supports Proliferation of Triple-negative Breast Cancer Cells and Contributes to mTORC1-S6K1 Pathway Activation. J Biol Chem 2014;289(6):3138–47 doi 10.1074/jbc.M113.502633.

29. Trauzold A, Schmiedel S, Sipos B, Wermann H, Westphal S, Roder C, et al. PKC mu prevents CD95-mediated apoptosis and enhances proliferation in pancreatic tumour cells. Oncogene 2003;22(55):8939–47 doi 10.1038/sj.onc.1207001.

30. Jaggi M, Rao PS, Smith DJ, Wheelock MJ, Johnson KR, Hemstreet GP, et al. E-cadherin phosphorylation by protein kinase D1/protein kinase C{mu} is associated with altered cellular aggregation and motility in prostate cancer. Cancer Res 2005;65(2):483–92.

31. Alpsoy A, Gunduz U. Protein kinase D2 silencing reduced motility of doxorubicin-resistant MCF7 cells. Tumor Biol 2015;36(6):4417–26 doi 10.1007/s13277-015-3081-3.

32. Ito Y. RUNX genes in development and cancer: Regulation of viral gene expression and the discovery of RUNX family genes. Adv Cancer Res 2008;99:33-+ doi 10.1016/S0065-230x(07)99002-8.

33. Pratap J, Lian JB, Javed A, Barnes GL, van Wijnen AJ, Stein JL, et al. Regulatory roles of Runx2 in metastatic tumor and cancer cell interactions with bone. Cancer metastasis reviews 2006;25(4):589–600 doi 10.1007/s10555-006-9032-0.

34. Komori T. Regulation of bone development and maintenance by Runx2. Front Biosci-Landmrk 2008;13:898–903 doi 10.2741/2730.

35. Borges S, Perez EA, Thompson EA, Radisky DC, Geiger XJ, Storz P. Effective Targeting of Estrogen Receptor-Negative Breast Cancers with the Protein Kinase D Inhibitor CRT0066101. Molecular cancer therapeutics 2015;14(6):1306–16 doi 10.1158/1535-7163.MCT-14-0945.

36. Harikumar KB, Kunnumakkara AB, Ochi N, Tong ZM, Deorukhkar A, Sung BY, et al. A Novel Small-Molecule Inhibitor of Protein Kinase D Blocks Pancreatic Cancer Growth In vitro and In vivo. Molecular cancer therapeutics 2010;9(5):1136–46 doi 10.1158/1535-7163.MCT-09-1145.

37. Bernhart E, Damm S, Wintersperger A, DeVaney T, Zimmer A, Raynham T, et al. Protein kinase D2 regulates migration and invasion of U87MG glioblastoma cells in vitro. Exp Cell Res 2013;319(13):2037–48 doi 10.1016/j.yexcr.2013.03.029.

38. Bernhart E, Damm S, Heffeter P, Wintersperger A, Asslaber M, Frank S, et al. Silencing of protein kinase D2 induces glioma cell senescence via p53-dependent and -independent pathways. Neuro-Oncology 2014;16(7):933–45 doi 10.1093/neuonc/not303.

