## Supplementary material for "Protein kinase D promotes prostate cancer cell bone metastasis by positively regulating Runx2 in a MEK/ERK1/2-dependent manner": FigS1 and Table S1

**Supplemental Materials**

A

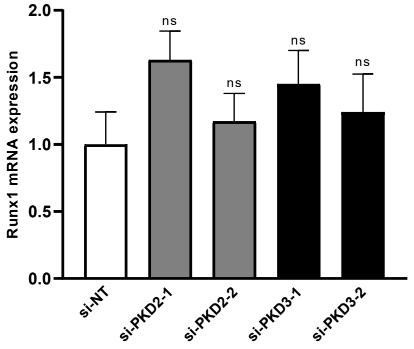


B

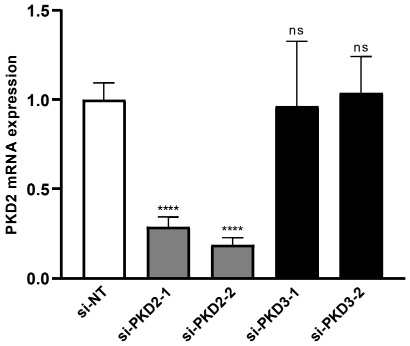


C

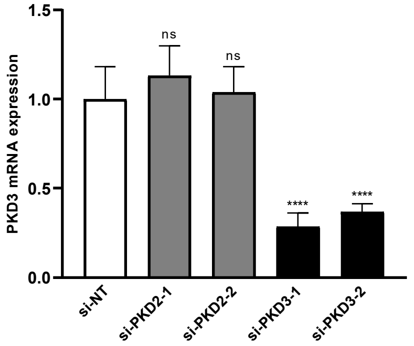


**Figure S1. The effect of PKD knockdown or inactivation on Runx1 and Runx2 transcription in prostate cancer cells.** (**A-C**) PC3-ML cells were transfected with PKD2 or PKD3 siRNA for 48 h or treated with CRT101 (5μM) for 24 h. RT-qPCR was performed for the Runx1 (A), PKD2 (B), and PKD3 (C) mRNAs. Transcript levels relative to the GAPDH control were determined. Data represent the Mean ± SEM from two independent experiments with triplicate determinations.

**Table S1. List of real time PCR primers used in the study.**

| Gene name | Forward primer (5′–3′) | Reverse primer (5′–3′) |
|---|---|---|
| Runx1 | AACCTCGAAGACATCGGCAG | GGCTGAGGGTTAAAGGCAGT |
| Runx2 | GAGTGGACGAGGCAAGAGTT | GGATGAGGAATGCGCCCTAA |
| Runx3 | CGGGGACCCTAACAACCTTC | GTGGGGGTGGTAACCTATGC |
| OPN | GCCCATCCCGTAAATGAAAAAG | GCTGACAACCAAGCCCTCCCAG |
| OCN | GCTTTGTTTACTTGTCAGGTTGGG | CCCTGTGTCCTTAGCAGGCAGGGA |
| MMP13 | TTTTGAGACCCTGCTGAAACAA | GTCTTTCCGCAGAGATTACC |
| FAK | GGTGCCTGGAGAGTGCAC | GCCGGGGTCGCCCCGCCGA |
| MMP9 | CCATCACTTTCCCTTGGCT | ACCAGCATGAGAAAGGGCTT |
